# Detecting the Information Flow in Proteins by Hodge Decomposition

**DOI:** 10.64898/2026.07.23.740308

**Authors:** Aysima Hacisuleyman

## Abstract

Allosteric communication in proteins is commonly quantified as a directed or undirected coupling between residues, but such descriptors mix distinct modes of signalling into a single pattern. Here we treat the net transfer entropy flux from a dynamic Gaussian network(dGNM) model as an edge flow on the residue contact graph and apply the combinatorial Hodge decomposition, which dissects the flow orthogonally into a gradient (global source-to-sink hierarchy), a curl (local three-clique circulation) and a harmonic (cavity-scale circulation) component. Applied to the wild-type KRAS and ten oncogenic KRAS variants spanning the principal GTPase-cycle mechanism classes, partial-hydrolysis position-12 (G12D, G12C, G12S), GAP-occluding position-12 (G12V, G12R), catalytic switch II (Q61R, Q61H), fast-cycling (G13D, A146T) and a combined steric and catalytic double mutant (G12D/Q61H), on a side-chain-centroid contact network, the decomposition shows that the transfer entropy flux is overwhelmingly hierarchical: the gradient term carries 97.5–98.4% of the flux in every variant (permutation p = 0.002), and the recovered scalar potential is strongly anti-correlated with each residue’s net outgoing transfer entropy (Spearman ρ ≈ −0.92 to −0.95). The hierarchy is conserved in magnitude but relocated by mutations: the dominant information sources move from the C-terminal α_5_/hypervariable region in wild type into the nucleotide-processing core, the switch I/II machinery and the α4/distal lobe in a way that tracks the GTPase-cycle mechanism of the substitution, while the sinks remain fixed. The method provides a parameter-free, residue-level readout of how mutations of different mechanism reposition the source of allosteric signalling in KRAS.

## Introduction

Allostery is the communication system in proteins where a perturbation at one residue propagates throug the packed residue networks to couple remote regions.^1-4^ A large amount of computational work has been made to assign a measure on how strongly and in which direction this dynamic information is communicated. Methods based on mutual information(MI), transfer entropy(TE) and network-flow centralities have all ben used to trace and uncover the commuication pathways between functional sites.^5-9^

Despite their utility, existing descriptors are often difficult to interpret because they compress several distinct modes of communication into a single directed coupling pattern.^2, 9^ In particular, a single matrix or edge-wise score may simultaneously reflect a global source-to-sink hierarchy, local recirculatory exchange, and larger-scale cycles spanning functionally relevant regions, without providing a canonical way to separate these contributions. This makes it hard to distinguish hierarchical signal propagation from genuinely circulatory communication, and to assign residue roles in a parameter-free way. The need for such a separation motivates a flow-based formulation in which directed allosteric communication is analyzed as an edge flow on a graph.^10^

We address these limitations by observing the directed allosteric communication in a protein structure as a flow field on a graph, which admits a classical and canonical decomposition. In the continuous setting, the Helmholtz-Hodge theorem splits any vector field into an irrotational (gradient) part, a solenoidal(curl) part and a harmonic part. Its analogue, combinatorial Hodge theory, performs the same orthogonal decomposition for a flow defined on the edges of a graph.^10, 11^ Applied to the net transfer entropy flux derived from an elastic network model (ENM), this decomposition yields directly the quantities that existing descriptors lack: a parameter-free ranking of residues, a clean separation of hierarchical communication from local ccirculatory communication and a topological observable, the harmonic component that quantifies the information circulating around proteins functional cavities.

Before the formal development it is useful to state what each component represents physically. The gradient part is a hierarchical, source-to-sink flow: much as water runs downhill across a landscape of altitudes, information moves from residues that sit high on a scalar potential, sources that broadcast signal, to those that sit low, sinks that receive it, with no closed loops. A single number per residue, its potential, therefore orders the whole hierarchy, and the flow along any contact is fixed by the difference in potential between its two residues. Curl flow, in contrast, is local: information passes around a triangle of mutually contacting residues and returns to where it started, a reciprocal exchange with no net source or sink. The harmonic part circulates on a larger scale still. Here information loops around a cavity or hole in the contact network without any local rotation and without a source or sink at all; the flow is purely topological, and whether it exists depends only on the shape of the network, that is, on how many independent loops it has. Since the three parts are mutually orthogonal, the decomposition separates the directed hierarchy cleanly from the two circulatory modes and assigns each a well defined share of the total flux.

The mathematical machinery is well established. The combinatorial Hodge decomposition, together with its use to rank inconsistent pairwise comparisons under the name HodgeRank was intoduced by Jiang et al.^10^ and Hodge-based decompoition have already been applied to neural networks.^12-17^ In structural biology, Hodge theory on simplical complexes has recently been used for biomolecular data analysis, persistent-spectral and Hodge-Laplacian descriptors of macromolecular structure by Koh et al.^18^, establishing that the Hodge Laplacian and its harmonic (Betti) spectrum carry biologically meaningful structural information. Our work brings this concept into protein allostery; we interpret the three components of the transfer entropy flux as hierarchical, local-circulatory and cavity-circulatory communication, and we equip the source-to-sink hierarchy with a permutation based significance test which makes its statistical significance assessable.

We demonstrate the method on the small GTPase KRAS, comparing its wild type with ten oncogenic variants that span the principal mechanistic classes of KRAS activation: the wild type (WT) (PDB id: 6GOD)^19^; the partial-hydrolysis position-12 substitutions^20^ G12D (PDB id: 6GOF)^19^, G12C (PDB id: 6OIM)^21^ and G12S (PDB id: 7TLK)^22^; the GAP-occluding position-12 substitutions^23^ G12V (PDB id: 8QDN)^24^ and G12R (PDB id: 6CU6)^25^; the catalytic switch II mutants Q61R (PDB id: 9O0S)^26^ and Q61H (PDB id: 6MNX)^27^; the fast-cycling mutants^28^ G13D (PDB id: 8EBZ)^29^ and A146T (PDB id: 8EDY)^30^; and the combined steric-plus-catalytic double mutant G12D/Q61H (PDB id: 9U8Z)^31^. KRAS is a paradigm allosteric switch whose activity is governed by communication between the nucleotide-binding P-loop and the switch I / switch II regions, and whose oncogenic mutations perturb this machinery. Across the WT and all of the ten variants we find that the source-to-sink hierarchy overwhelmingly dominates the total transfer entropy flux, while the small circulatory (curl and harmonic) components and the number of independent contact loops shift with the mutation, and the emitting end of the hierarchy migrates from the C-terminal α_5_/hypervariable region in wild type into the nucleotide-processing core, and which core region receives it reflects the GTPase-cycle mechanism of each substitution.

## Methods

### Structures and elastic-network representation

Each protein is represented as a graph whose nodes are residues placed at the centroid of their side-chain heavy atoms and whose edges join residues in spatial contact. For each residue the contact coordinate is the mean position of all side-chain heavy atoms (backbone N, Cα, C, O, with hydrogens excluded); glycine, which has no side chain, is assigned a virtual Cβ reconstructed from its backbone N, Cα and C atoms. Atomic coordinates were taken from the crystal structures listed above. A residue pair is connected when the distance between their side-chain centroids is below a cutoff r_c_ (7.5⍰Å for KRAS). The resulting contact Kirchhoff matrix, Γ, has Γ_*ij*_ = −1 for contacting pairs and Γ_*ii*_ = − ∑_*i*≠*j*_ Γ_*ij*_ on the diagonal.

### Transfer entropy flux

We quantify the directed allosteric communication by the transfer entropy *T*_*i*→*j*_ computed from the dynamic Gaussian Network Model(dGNM) presented by Hacisuleyman and Erman^32^ from Schreiber’s^33^ entropy transfer concept, which measures how much the past fluctuations of residue *i* reduce the uncertainty about the future fluctuations of residue *j*. From the directional pair *T*_*i*→*j*_ and *T*_*j*→*i*_ we construct the *net* information flux on each edge,

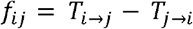

an antisymmetric quantity (*f*_*ij*_ = −*f*_*ji*_) which is positive when residue *i* is the information source for residue *j*. The collection of all *f*_*ij*_ defines a single discrete flow field on the contact graph. The aim is to decompose this field into interpretable components. The transfer entropy is evaluated at a single characteristic time delay τ, chosen automatically from the GNM dynamics rather than set by hand: for each residue we compute the normalised autocorrelation of its position fluctuations, 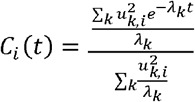 (sum over non-zero GNM modes *k* with an eigenvalue *λ*_*k*_ and an eigenvector component *u*_*k,i*_), and define its relaxation time as the first time *t* where *C*_*i*_*(t)* decays to 1/e of its initial value. The single delay used for the whole protein is the median of these per-residue relaxation times, τ = median_i_ τ_i_, which gives τ = 0.33 (WT), 0.31 (G12D), 0.32 (G12C), 0.31 (G12S), 0.37 (G12V), 0.33 (G12R), 0.31 (Q61R), 0.29 (Q61H), 0.33 (G13D), 0.31 (A146T) and 0.31 (G12D/Q61H) GNM time units. Using the median makes the delay a robust, parameter-free property of each structure’s own dynamics.

A raw matrix of directed couplings combines three physically distinct phenomena: a global source-to-sink hierarchy, information that recirculates locally, and information that circulates around functional cavities. Combinatorial Hodge theory, the discrete analogue of the Helmholtz decomposition of vector fields, separates any edge flow into exactly these three parts. Because they are mutually orthogonal, every unit of information flux belongs unambiguously to one mechanism, and we can measure how much of the communication each one carries.

### The Hodge decomposition

Beyond nodes and edges we include the graph’s 3-cliques (triples of mutually contacting residues) as triangular 2-simplices. Two linear operators act on this complex. The gradient operator *G* maps a scalar defined on residues, (*Gϕ*)_*ij*_ = *ϕ*_*j*_ − *ϕ*_*i*_ ; intuitively it turns a “height” assigned to each residue into a downhill flow. The curl operator *C* maps an edge flow to a value on each triangle, (*Cf*)_*ijk*_ = *f*_*ij*_ + *f*_*jk*_ + *f*_*ki*_, *mea*suring the net circulation around that triangle. These operators satisfy the fundamental identity *CG* = 0: a pure downhill flow has no circulation.

The space of edge flows splits orthogonally into the image of the gradient, the image of the curl adjoint, and the kernel of the 1-Hodge-Laplacian *L*_1_ = *GG*^*T*^ + *C*^*T*^*C*. Any net flux therefore decomposes uniquely as

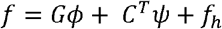

Where *Gϕ* is the gradient, *C*^*T*^*ψ* is the curl and *f*_*h*_ is the harmonic components. Since the three subspaces are orthogonal the squared magnitudes add, ∥*f*∥^2^ = ∥*Gϕ*∥^2^ + ∥*C*^*T*^*ψ* ∥^2^ + ∥*f*_*h*_ ∥^2^. The fraction of the total in each term is reported as the gradient, curl, and harmonic share of the communication. We obtain the components by two least-squares projections: the residue potential *ϕ* solves the graph-Laplacian system *L*_0_*ϕ = G*^*T*^*f* (with *L*_0_ *= G*^*T*^*G* the ordinary Kirchhoff Laplacian), the curl potential *ψ* solves *CC*^*T*^*ψ = Cf*, and the harmonic part is the remainder *f*_*h*_ *=f − Gϕ − C*^*T*^*ψ*. We solve both systems with the minimum-norm pseudoinverse, since *L*_0_, is rank-deficient by one (the additive constant in *ϕ*).

### Interpretation of the three components

The *gradient* component is the irrotational, source-to-sink part. The scalar potential *ϕ* recovered, up to an additive constant, by least-squares fitting of a residue “height” to the observed flux, provides a parameter-free ranking of every residue along the allosteric hierarchy, requiring no a priori choice of source or sink. Residues at the low extreme of *ϕ* are net information emitters; those at the high extreme are net receivers. This is the discrete analogue of HodgeRank, the same construction used to extract a globally consistent ranking from inconsistent pairwise comparisons.^10^

The *curl* component is the locally circulating part, supported on filled triangles. It represents information that cycles among three mutually contacting residues and cannot be explained by any global hierarchy.

The *harmonic* component is the part that is simultaneously source-free and curl-free yet still circulates. By the discrete Hodge theorem, the dimension of the harmonic subspace equals the first Betti number *β*_1_ of the complex, the number of independent loops not bounded by triangles, i.e. the topological holes of the contact network.^18^ Harmonic flux can only circulate around such a hole, so a non-zero harmonic component identifies information looping around a functional cavity (for example, a ligand-binding cleft). A vanishing harmonic component indicates purely hierarchical signaling.

### Choice of graph substrate: contact vs. top-flux edges

The decomposition is defined relative to a graph, and the interpretation of the harmonic component rests on the graph faithfully representing the protein’s geometry, because that interpretation requires the loops to be real topological holes. We therefore compute the full three-way decomposition, including the harmonic component and β_1_, exclusively on the residue contact graph (“contact” mode), whose cycles correspond to genuine cavities and clefts in the fold. We do not use the top-flux (k-nearest-neighbour, “tetop”) graph, in which each residue is joined to its k strongest net-flux partners, for the harmonic analysis: the loops of such a graph are artefacts of the kNN construction rather than real holes, so its Betti numbers and harmonic flux are not physically interpretable. The gradient (source-to-sink) ranking, by contrast, is robust to the choice of substrate. Where a robustness check is desired, we therefore verify that the residue ranking derived from the potential *ϕ* is preserved when the contact graph is replaced by the top-flux graph; consistency of the ranking across the two substrates is used as a control, while all reported harmonic and Betti results are computed on the contact graph. All results below were obtained in contact mode.

### Significance of the hierarchy

To assess whether the observed hierarchy exceeds chance, we compare the gradient share of the flux against a null distribution generated by randomly permuting the net-flux values across edges and recomputing the decomposition (500 permutations). The reported p-value is the fraction of permutations whose gradient share equals or exceeds the observed value. As an internal consistency check we also report the Spearman correlation between the recovered potential *ϕ* and the net outgoing transfer entropy of each residue, which is expected to be strongly negative (low *ϕ* = source = high net-out TE).

## Results

### The transfer entropy flux is overwhelmingly hierarchical in every KRAS variant

Across all of the eleven structures the Hodge decomposition assigns the great majority of the transfer component: 98.4% in wild-type KRAS (6GOD) and between 97.5% and 98.4% across the oncogenic variants (Table 1, Fig. 1a). The curl (local 3-clique) and harmonic (cavity-loop) components together carry under 3% of the flux in every case, curl 1.0–1.7% and harmonic 0.4–1.1% (Fig. 1b). The dominance of the gradient term is highly significant: in every variant the permutation test returns p = 0.0020 (500 permutations), with an observed gradient share of 0.975–0.984 against a null mean of only ∼0.2–0.3. The recovered potential *ϕ* is strongly anti-correlated with each residue’s net outgoing transfer entropy (Spearman ρ between −0.919 and −0.949), confirming that low-*ϕ* residues are the information sources as intended. KRAS allosteric communication is therefore predominantly a directed, hierarchical signal rather than a circulatory one.

**Table 1.**
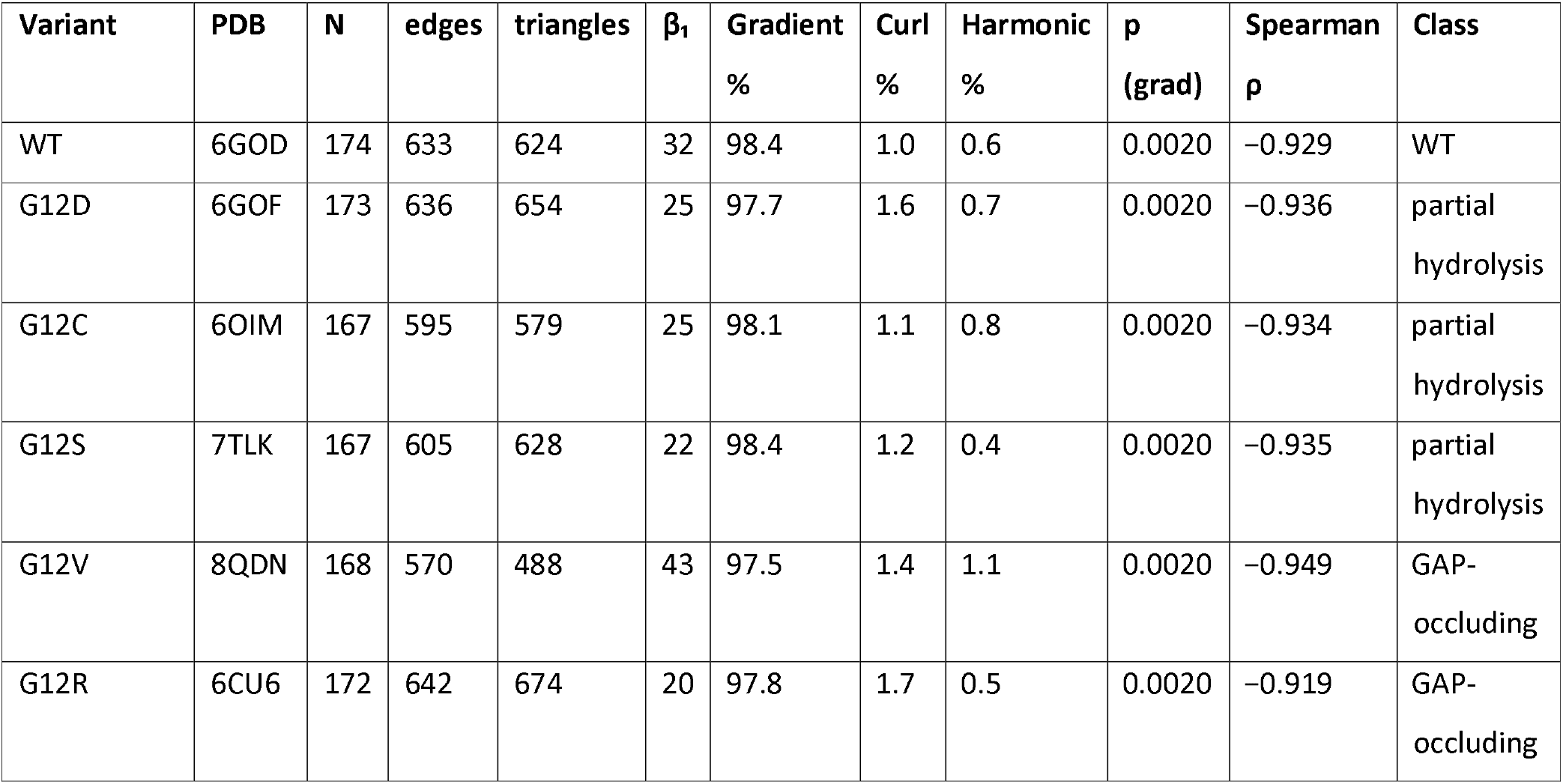
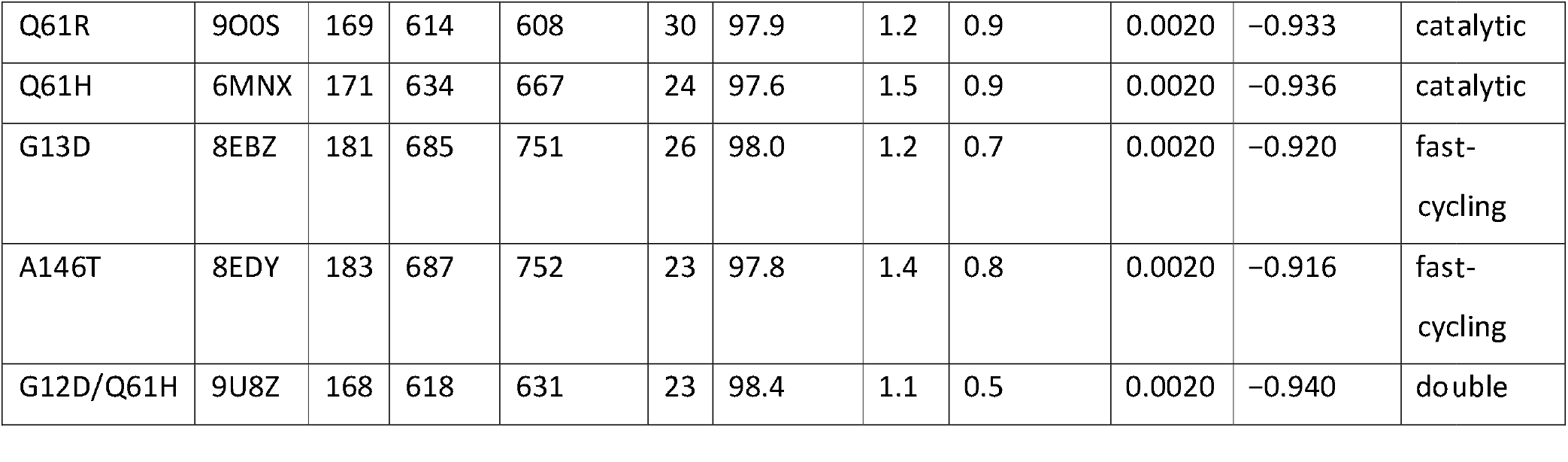
Hodge decomposition of the GNM transfer entropy flux for the eleven KRAS variants on the side-chain-centroid contact graph (7.5 Å cutoff, virtual Cβ for glycine). N, number of residues; edges and triangles, the 1- and 2-simplices of the contact complex; β_1_, first Betti number; Gradient/Curl/Harmonic %, share of the total flux carried by each Hodge component; p (grad), permutation p-value for the gradient share (500 permutations); Spearman ρ, correlation between the recovered potential *ϕ* and each residue’s net-out transfer entropy; Class, GTPase-cycle mechanism class. Variants are grouped by mechanism class.

**Figure 1.**
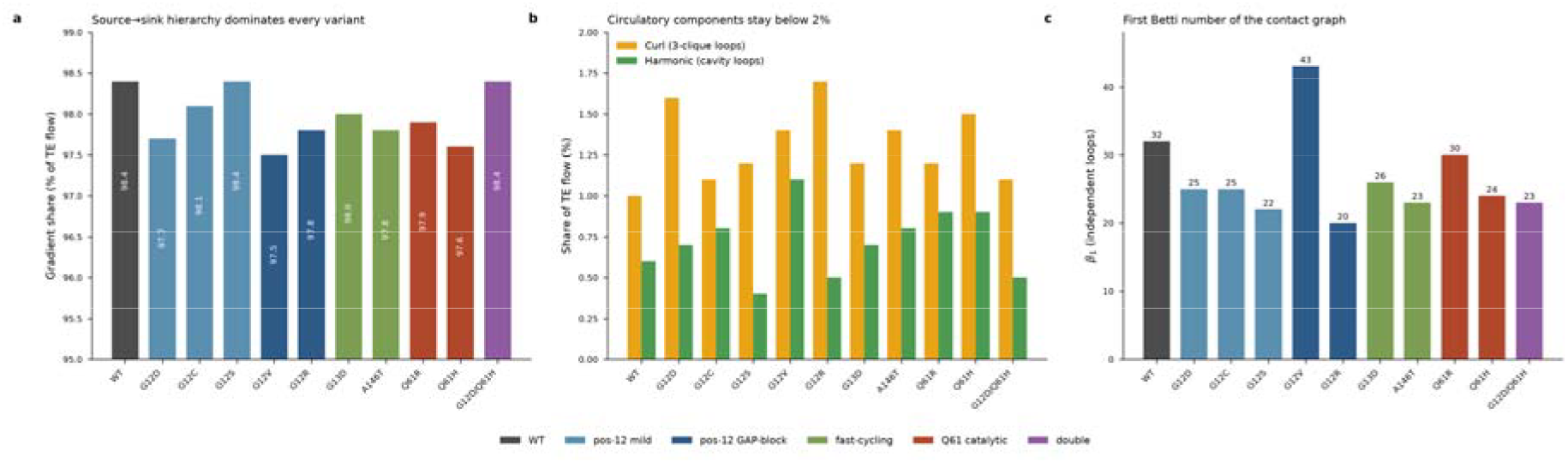
Hodge decomposition of the transfer entropy flux across the eleven KRAS variants, ordered by GTPase-cycle mechanism class (bar colour). (a) The gradient (source-to-sink) component carries 97.5–98.4% of the flux in every variant (note the truncated 95–99% axis). (b) The curl (local 3-clique) and harmonic (cavity-loop) components together account for under 2% of the flux. (c) The first Betti number β_1_ of the side-chain contact graph, the number of independent loops available to harmonic circulation, varies with the mutation (range 20–43). All quantities were computed in contact mode on the side-chain-centroid graph (7.5 Å cutoff, virtual Cβ for glycine); the permutation test gives p = 0.002 for the gradient share in every variant.

### The emitting source relocates from the C-terminal lobe into the nucleotide-processing core along GTPase-cycle mechanism class

In wild-type KRAS the strongest information sources (most negative ⍰, highest net-out TE) are the C-terminal α_5_/hypervariable residues (168, 171, 172) together with the α4/distal-lobe residues 105 and 121; the sinks lie in the N-terminal/β_1_ region and around the β_4_–α3 lobe and are conserved across every variant. What the mutation changes is not the strength of the hierarchy but where its emitting end sits, and that shift follows the substitution’s place in the GTPase cycle. In wild type the C-terminal α_5_/hypervariable region alone supplies 63% of the total top-source emitting strength; in every oncogenic variant this share collapses to 27% or below and the emitting strength is redistributed into the nucleotide-processing core. The partial-hydrolysis position-12 substitutions (G12D, G12C, G12S) move the leading emitters onto switch I (residue 31) and switch II (residues 63– 67), raising the combined switch I + II share from 6% in wild type to 19–72%. The GAP-occluding position-12 substitutions G12V and G12R carry the largest switch region loads: switch II residues 65– 70 dominate G12V while switch I residues 30–31 lead G12R, with the α4/distal lobe (residues 105, 121) providing a secondary source in both. The catalytic switch II mutants shift the emitting end predominantly into the α4/distal lobe: residues 105, 121 and 122 lead Q61R and residues 105, 121, 128 and 132 lead Q61H, with a smaller direct switch II contribution (residue 73 in Q61R). The fast-cycling mutant G13D retains a mixed profile, with switch I (residues 30, 32, 34) and the α4/distal lobe (residue 105) sharing the load alongside a residual C-terminal contribution. The other fast-cycling variant, A146T (8EDY), instead concentrates its emitting strength almost entirely on switch I: eight of its ten leading sources fall in residues 32–39, with a single switch II contribution (residue 64), giving a switch I share near 73% and a Spearman ρ of −0.916. The combined steric-plus-catalytic double mutant G12D/Q61H superposes both signatures, drawing sources from switch I (residue 31) and from an extended α4/distal-lobe segment (residues 105, 128, 132, 135). The overall picture (Fig. 3) is that the emitting source migrates away from the C-terminal α_5_/hypervariable region in a mechanism-class-dependent way: the position-12 mutants load the switch regions directly, whereas the catalytic switch II mutants project their sources onto the α4/distal lobe that packs against switch II.

**Figure 2.**
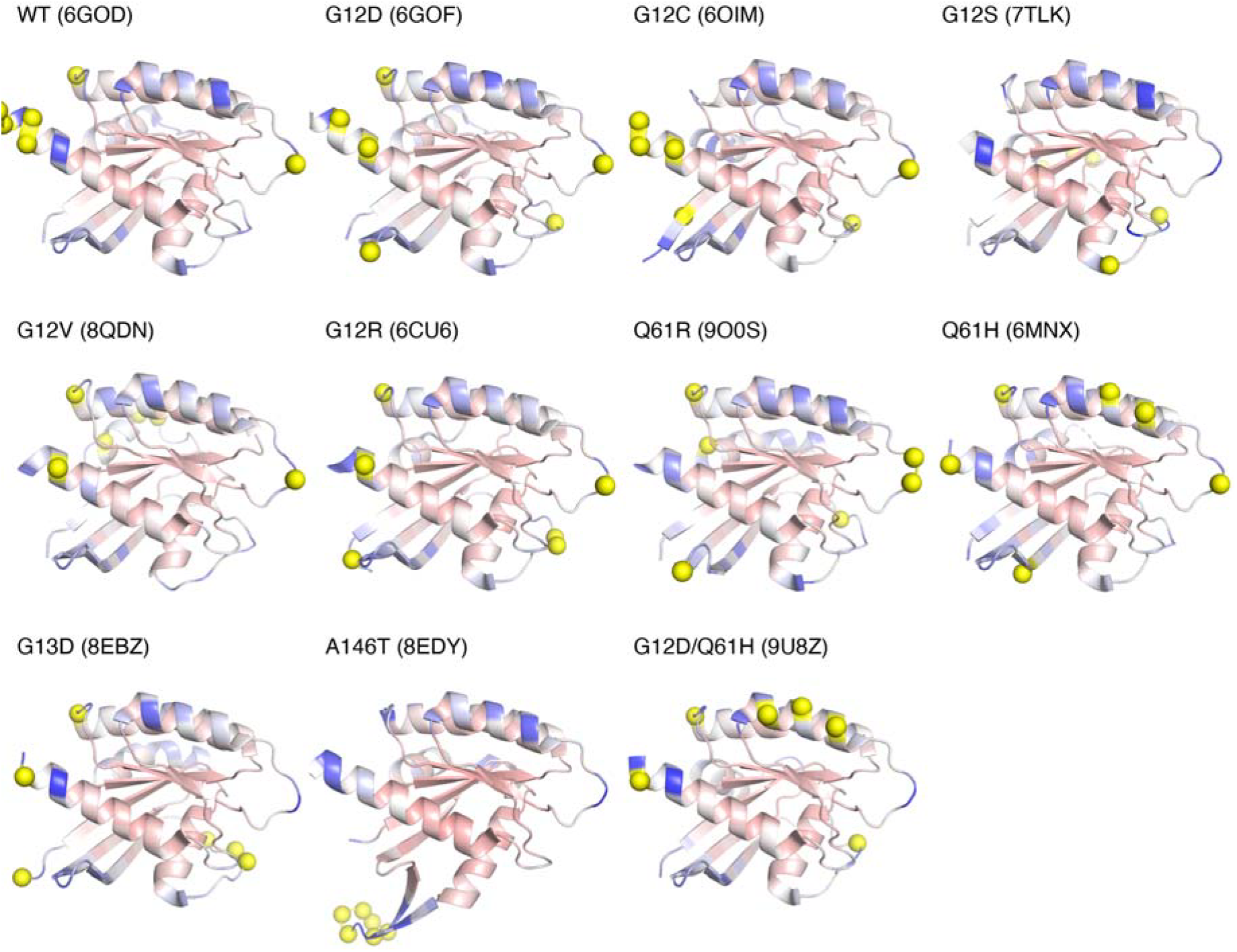
Scalar potential ⍰ mapped onto the KRAS structure for the wild type and all ten oncogenic variants, ordered by GTPase-cycle mechanism class. In every panel the cartoon is coloured by the Hodge scalar potential ⍰ on a shared blue–white–red scale (blue = information source, low *ϕ*; red = sink, high *ϕ*), and the six strongest information sources are drawn as yellow spheres. All structures are superposed on the WT frame. The source spheres migrate from the C-terminal α_5_/hypervariable lobe in the wild type toward the switch I/switch II region and the α4/distal lobe in the oncogenic variants, in a manner that tracks the mechanism class of the substitution, while the overall ⍰ field and the sink regions are conserved. A146T (8EDY) is the extreme case, concentrating its emitting sources almost entirely on switch I (lower-left cluster of spheres).

**Figure 3.**
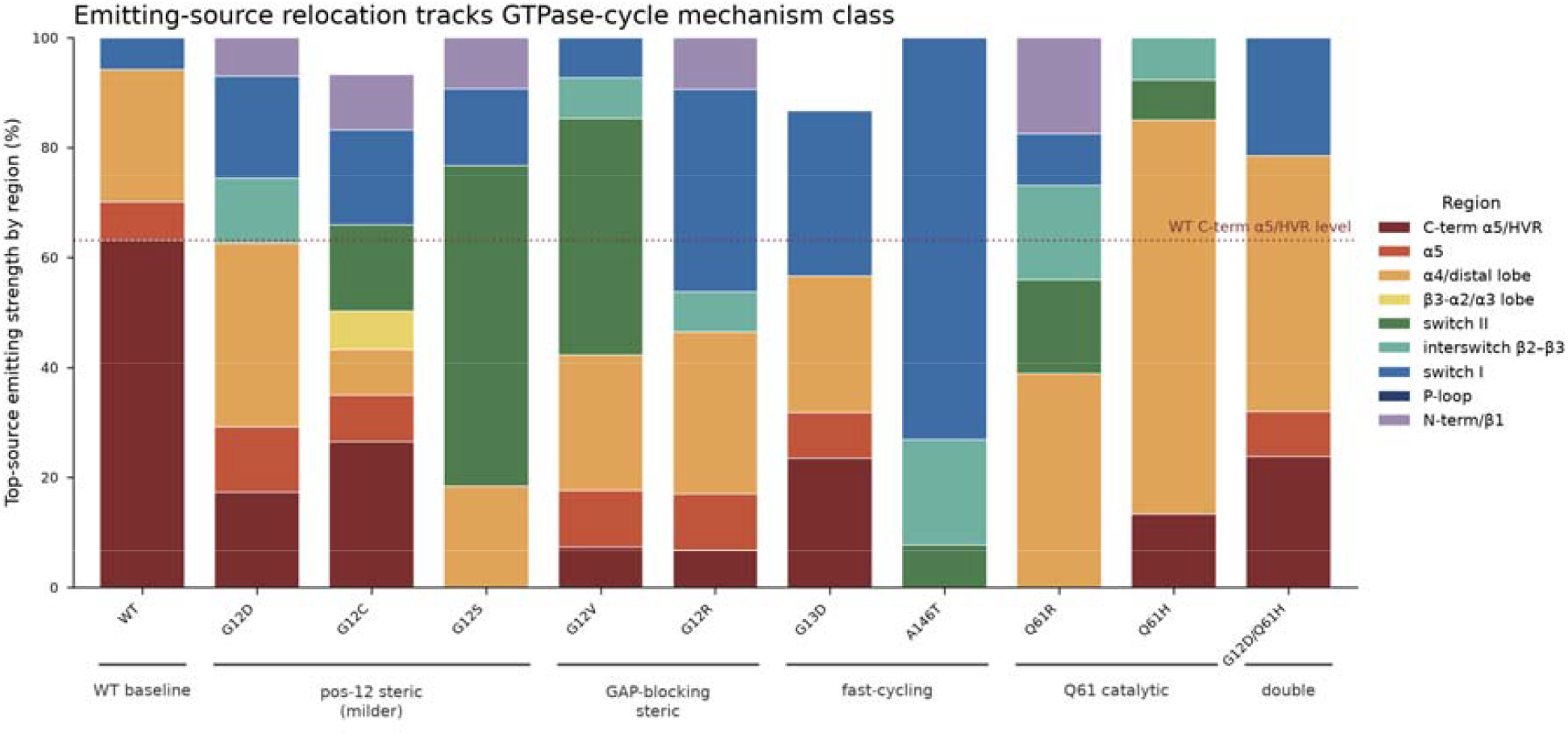
Emitting-source relocation by GTPase-cycle mechanism class. Emitting strength of each residue is defined as max(0, −), the positive part of the negative scalar potential, and the top-ten sources of each variant are grouped into functional regions and expressed as a percentage of that variant’s total top-source emitting strength. Variants are ordered and bracketed by mechanism class. The C-terminal α_5_/hypervariable region (dark red) carries 63% of the emitting strength in wild type (dotted reference line) but ≤27% in every mutant; the partial-hydrolysis and GAP-occluding position-12 substitutions load switch I/II directly, whereas the catalytic switch II mutants (Q61R, Q61H) project onto the α4/distal lobe. Because the scalar potential is steeply ranked, the ten leading sources capture nearly all of the negative- (emitting) mass, so the regional fractions are insensitive to the exact cutoff. Functional regions: N-terminal/β_1_ 1–9 and 18–29, P-loop 10–17, switch I 30–38, interswitch β_2_–β_3_ 39–56, switch II 57–76, β_3_–α2/α3 lobe 77–104, α4/distal lobe 105–151, α_5_ 152– 166, C-terminal α_5_/HVR 167–189.

### Mutations reshape the contact-network topology and relocate the harmonic circulation

The first Betti number of the contact graph, the count of independent loops available to carry harmonic flux, reorders with the mutation and spans β_1_ = 20–43 across the eleven variants (WT 32; G12D 25, G12C 25, G12S 22; G12V 43, G12R 20; Q61R 30, Q61H 24; G13D 26, A146T 23; G12D/Q61H 23; Table 1, Fig. 1c). The GAP-occluding G12V carries by far the most independent loops (43), while the other GAP-occluding substitution G12R carries the fewest (20), so loop count does not by itself order the variants by severity. Although the harmonic share of the flux is small in absolute terms (0.4–1.1%), the residues that carry it move with the mutation. In wild type and G12D the harmonic circulation is concentrated in the interswitch β_2_–β_3_ region and the α_5_ helix; in G12C it additionally invades the P-loop, the α_3_ helix and switch II, and in the loop-rich G12V it spreads away from the switches into the α4/distal lobe while retaining an interswitch contribution. The harmonic component thus provides a compact, topology-aware readout of how each oncogenic substitution redistributes the small circulatory portion of the communication.

## Discussion

The side-chain-centroid contact graph resolves a mutation-dependent reorganisation of the allosteric hierarchy across a panel of eleven KRAS structures. In every variant the transfer entropy flux is overwhelmingly gradient-like (97.5–98.4%), so allosteric communication is predominantly a directed source-to-sink process rather than a circulatory one. What changes with the oncogenic substitution is not the magnitude of this hierarchy but its geography, and the direction of that change follows the GTPase-cycle mechanism by which each mutation drives KRAS activation.^34^ In wild type the dominant information sources are the C-terminal α_5_/hypervariable residues, which alone carry 63% of the emitting strength; in every mutant this C-terminal dominance collapses and the emitting end migrates into the nucleotide-processing core, but where it lands depends on the class of mutation.

The partial-hydrolysis position-12 substitutions (G12D, G12C, G12S), which retain measurable intrinsic GTP hydrolysis and are only partially GAP-resistant, shift the sources directly onto switch I and switch II. The bulkier GAP-occluding position-12 substitutions G12V and G12R, which sterically occlude the arginine finger and most strongly suppress GAP-mediated GTP hydrolysis,^35^ carry the heaviest switch region source loads. The catalytic switch II mutants Q61R and Q61H, which disable the intrinsic hydrolysis machinery around Gln61,^34^ instead project their sources onto the α4/distal lobe that packs against switch II, consistent with a perturbation centred on the catalytic apparatus rather than on the P-loop. The fast-cycling mutants G13D and A146T, which accelerate nucleotide exchange rather than blocking hydrolysis,^36^ differ in their source geography: G13D retains a mixed, partly wild-type-like distribution spread across switch I, the α4/distal lobe and a residual C-terminal contribution, whereas A146T concentrates almost three-quarters of its emitting strength on switch I. Both rank among the least transforming of the classes considered. Finally, the combined steric-plus-catalytic double mutant G12D/Q61H superposes the position-12 and Q61 signatures, drawing sources from both switch I and an extended α4/distal-lobe segment. Read together, the emitting-source geography recovered by the Hodge potential thus orders the variants not by a single scalar severity but by mechanism class, providing a parameter-free, residue-level readout of where each mutation repositions the source of allosteric signalling. The harmonic component, though small, offers a complementary topology-aware fingerprint: its supporting loops move from the interswitch/α_5_ region in wild type toward the P-loop, α_3_ and switch II in G12C, and disperse into the α4/distal lobe in the loop-rich G12V. Independent structural evidence reinforces this localisation: the engineered monobody 12D4, which selectively recognises oncogenic KRAS(G12D), achieves its recognition through a conserved hydrophobic network that docks onto switch II and the α3-helix^37^, the same switch II surface onto which the Hodge potential relocates the emitting sources of the position-12 and catalytic variants. The structural potential panels (Fig. 2) map the full per-residue *ϕ* field for all eleven variants, and the emitting-source quantification (Fig. 3) uses the top-ten ranked sources for the same set.

Methodologically, the decomposition is agnostic to the origin of the flow: any directed coupling defined on a residue graph, transfer entropy from all-atom molecular dynamics, mutual-information or dynamical-network couplings, or fluxes from Markov state models, can be decomposed in the same way, so the present elastic-network analysis is a linear, equilibrium special case of a broader family. Replacing the harmonic GNM flux with transfer entropy estimated from molecular-dynamics trajectories would capture the anharmonic, nucleotide-state-dependent coupling that a single-structure elastic network cannot, and would let the contact graph evolve along a trajectory rather than remain fixed. The construction also invites application well beyond KRAS: because the relocation of the emitting source reports the biochemical mechanism of a substitution, the same readout could help interpret variants of unknown significance in other switch GTPases and signalling proteins, and, more generally, flag the switch surfaces and cavity-bounding loops highlighted by the gradient and harmonic components as candidate sites for allosteric intervention. The principal limitations to address are the fixed contact-graph cutoff, which we set at 7.5 Å on side-chain centroids without retuning to a target mean degree, and the restriction of the decomposition to cliques up to triangles; a systematic calibration of graph construction, together with the inclusion of higher-order simplices, would sharpen both the curl and harmonic estimates. Even in its present linear form, the method converts a high-dimensional flux matrix into a single, statistically testable ranking of residues, and we expect this combination of interpretability and significance testing to make it increasingly useful as the underlying models of protein dynamics grow in richness.

## Data availability

All source code and data preprocessing scripts are publicly available at https://github.com/Aisima/Hodge_decomposition_TE.

